# Cyclic lipopeptides as membrane fusion inhibitors against SARS-CoV-2: new tricks for old dogs

**DOI:** 10.1101/2022.12.05.519140

**Authors:** Egor V. Shekunov, Polina D. Zlodeeva, Svetlana S. Efimova, Anna A. Muryleva, Vladimir V. Zarubaev, Alexander V. Slita, Olga S. Ostroumova

**Author notes:** These authors contributed equally to this work (Egor V. Shekunov, Polina D. Zlodeeva, Svetlana S. Efimova).

## Abstract

With the resurgence of the coronavirus pandemic, the repositioning of FDA-approved drugs against coronovirus and finding alternative strategies for antiviral therapy are both important. We previously identified the viral lipid envelope as a potential target for the prevention and treatment of SARS-CoV-2 infection with plant alkaloids [1]. Here, we investigated the effects of eleven cyclic lipopeptides (CLPs), including well-known antifungal and antibacterial compounds, on the liposome fusion triggered by calcium, polyethylene glycol 8000, and a fragment of SARS-CoV-2 fusion peptide (816-827) by calcein release assays. Differential scanning microcalorimetry of the gel-to-liquid-crystalline and lamellar-to-inverted hexagonal phase transitions and confocal fluorescence microscopy demonstrated the relation of the fusion inhibitory effects of CLPs to alterations in lipid packing, membrane curvature stress and domain organization. The effects of the compounds were evaluated in an *in vitro Vero*-based cell model, and aculeacin A, anidulafugin, iturin A, and mycosubtilin attenuated the cytopathogenicity of SARS-CoV-2 without specific toxicity.

## Introduction

By the beginning of autumn 2022, the World Health Organization [2] reported approximately 600 000 000 confirmed cases of COVID-19 and more than 6 000 000 deaths. The search for new antiviral drugs and approaches remains a high priority.

Infection by various enveloped viruses, including severe acute respiratory syndrome coronavirus 2 (SARS-CoV-2), requires fusion between the viral lipid envelope and the host plasma or endosomal membrane. The process of fusion of two membranes takes place in several successive stages with the formation of certain lipid intermediates [3–5]. In the last decade, the idea has been circulating in the literature that using active small molecules to change the physical properties of the lipid matrix of fusing membranes can prevent virus entry and infection [6,7]. An effective fusion inhibitor aimed at impeding such membrane fusion may emerge as a broad-spectrum antiviral agent. From this point of view, antimicrobial lipopeptides (usually composed of peptide heads and fatty acid tails) should be of great interest due to their amphiphilic nature and conical shape. The promotion of positive curvature stress by lipopeptides is believed to inhibit the formation of fusion intermediates with negative curvature. In particular, Yuan et al. [8] found that a cyclic lipopeptide from *Bacillus subtilis*, surfactin, can suppress the proliferation of porcine epidemic diarrhea virus and transmissible gastroenteritis virus in epithelial cells in a relatively low concentration range (15 to 50 μg/ml), without cytotoxicity or viral membrane disruption. A simple synthetic lipopeptide myr-WD has been shown to successfully combat type 1 influenza virus (H1N1) and murine coronavirus infections [9]. The authors of both cited works associated the antiviral effect of the lipopeptides with the suppression of the fusion of the viral and cell membranes by altering their properties.

Here, we studied the effects of 11 cyclic lipopeptides (CLPs) on membrane fusion triggered by calcium, polyethylene glycol 8000, and a short fragment of SARS-CoV-2 fusion peptide 1 (816-SFIEDLLFNKVT-827). The choice of CLPs was determined by proven wide-spectrum pharmacological activity against bacteria and fungi; in particular, well-known drugs such as polymyxin B, daptomycin, and echinocandins (caspofungin and anidulafungin) were tested. Changes in the elastic properties of membranes associated with the introduction of CLPs were assessed using differential scanning microcalorimetry and confocal fluorescence microscopy. We demonstrated that the inhibitory effect of CLPs on membrane fusion is related to the disordering of membrane lipids, the induction of positive curvature stress and enhancing raft formation. Finally, we found that aculeacin A, anidulafugin, iturin A, and mycosubtilin act as potent membrane fusion inhibitors combating SARS-CoV-2 cytopathogenicity.

## Results and Discussion

### CLPs are able to inhibit liposome fusion

We analyzed the effects of aculeacin A, anidulafungin, caspofungin, fengycin, iturin A, mycosubtilin, surfactin, syringostatin A, syringotoxin B, daptomycin, and polymyxin B, on the fusion of membranes of various compositions triggered by calcium, polyethylene glycol 8000 (PEG-8000), and a short fragment of SARS-CoV-2 fusion peptide 1 (816-SFIEDLLFNKVT-827, FP_SARS(816-827)_). A schematic representation of the structures of the CLPs used is shown in Figure 1. All compounds are characterized by an amphiphilic structure composed of a cyclic peptide “head” (containing 6 to 10 amino acid residues) and a hydrocarbon “tail” (8 to 18 carbon atoms successively connected in a chain).

**Figure 1.**
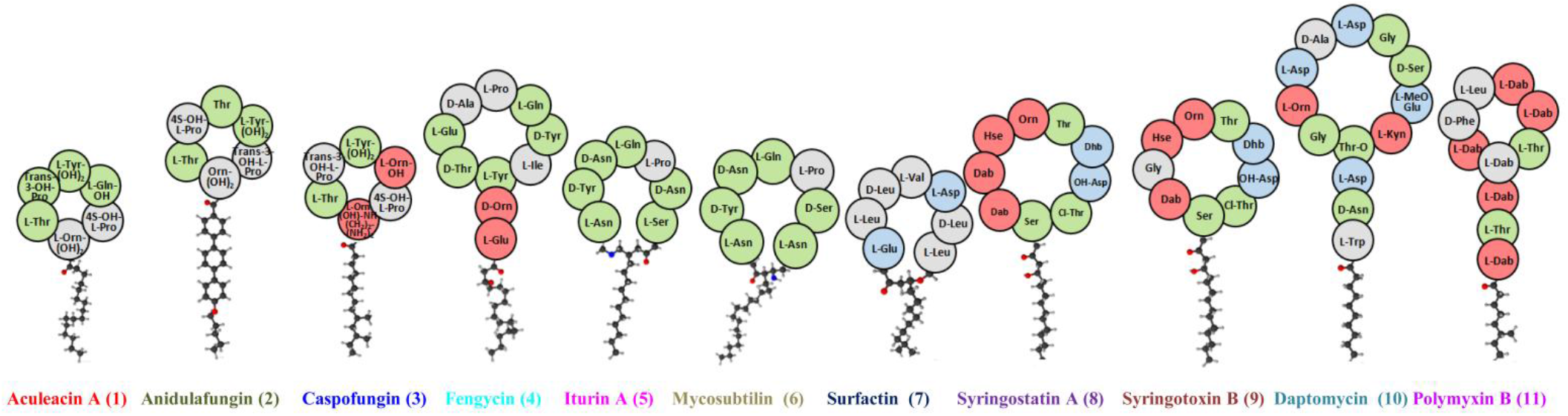
Schematic representation of the chemical structures of the tested CLPs. Information about CLP structures is taken from [10–18]. The red circles indicate positively charged amino acid residues, blue indicates negatively charged amino acid residues, green indicates neutral polar amino acid residues, and gray indicates neutral hydrophobic amino acid residues. Noncanonical amino acids: Dab, 2,4-diaminobutyric acid; Dhb, 2,3-dehydroaminobutyric acid; Hse, homoserine; Orn, ornithine; Kyn, kynurenic acid.

Figure 2a (upper panel) shows the time dependence of the percentage of fusion (*% fusion*) of negatively charged vesicles triggered by the calcium cations in the absence (black curve) and presence of the tested CLPs (curves of appropriate colors). The liposomes were composed of DOPC/DOPG/CHOL (40/40/20 mol.%). Control samples were not modified with CLPs. CLPs were added to the experimental samples and incubated with vesicles for 35 ± 5 minutes. The decrease in *% fusion* of liposomes pretreated with CLPs indicates the ability of the tested compounds to inhibit the calcium-induced fusion of the negatively charged vesicles. To compare the antifusogenic ability of different CLPs, we applied the maximum value of *% fusion* in the presence of appropriate CLPs averaged over several independent experiments. The values are summarized in Supplementary Table S2. The value of *% fusion* in the presence of CLPs decreased in the series: syringostatin A > polymyxin B > syringotoxin B ≥ iturin A > aculeacin A ≈ mycosubtilin > anidulafungin ≈ caspofungin ≈ fengycin. Surfactin and daptomycin were unable to suppress the calcium-mediated fusion of DOPC/DOPG/CHOL liposomes (Supplementary Table S2).

**Figure 2.**
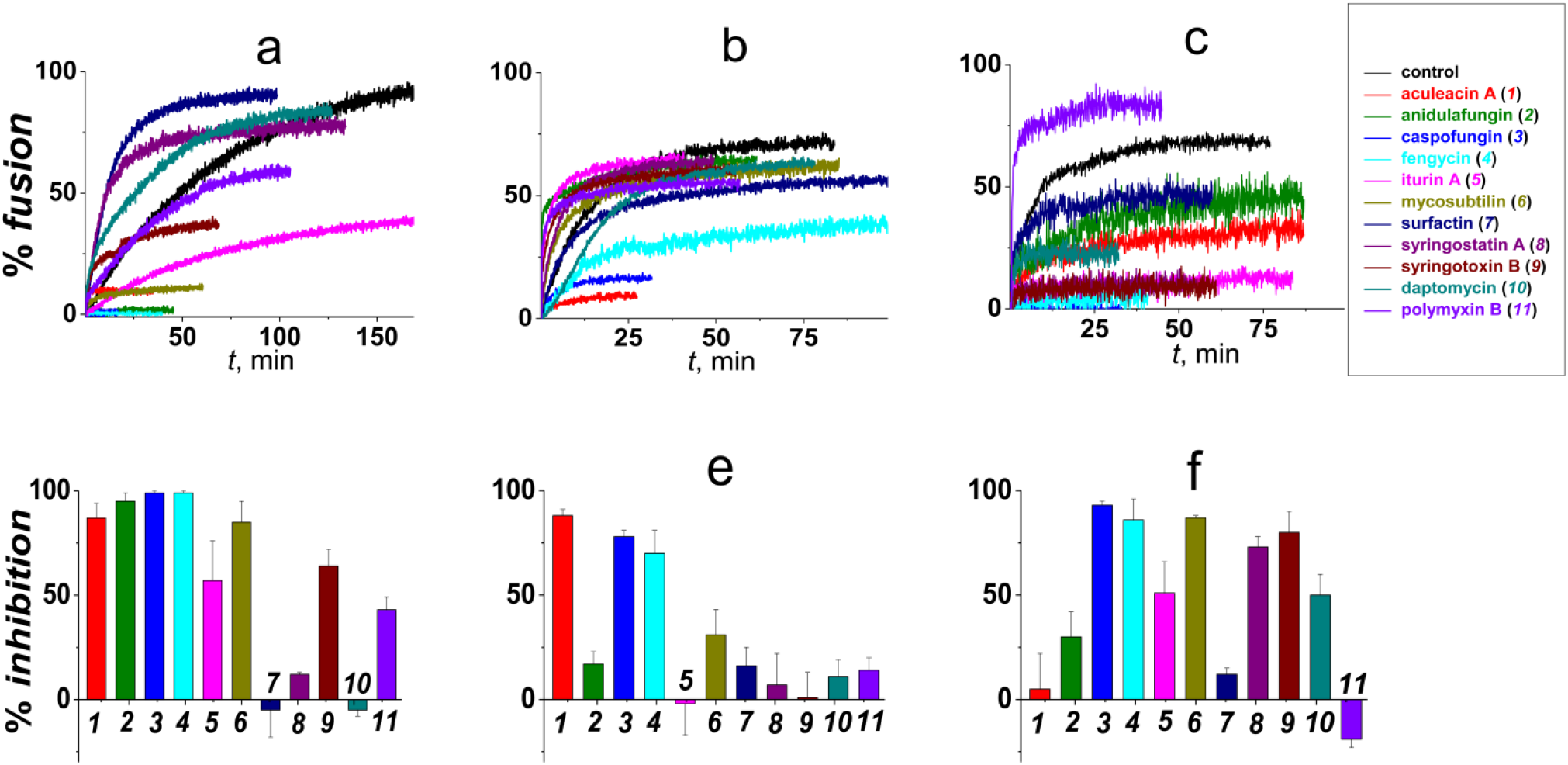
***Upper panel*** – Time dependence of the percentage of fusion (*% fusion*) of liposomes composed of DOPC/DOPG/CHOL (40/40/20 mol.%) (**a**), DOPC/CHOL (80/20 mol.%) (**b**), and POPC/SM/CHOL (60/20/20 mol.%) (**c**) mediated by 40 mM CaCl_2_, 20 wt.% PEG-8000, 150 μm FP_SARS(816-827)_, respectively. The relationship between the color and the CLP is shown in (**c**). Control samples were not modified with CLPs. *% fusion* was calculated using a calcein leakage assay. CLP concentrations are presented in Supplementary Table S1. ***Lower panel*** – The mean percentage of inhibition of fusion (*% inhibition*) of liposomes composed of DOPC/DOPG/CHOL (40/40/20 mol.%) (**a**), DOPC/CHOL (80/20 mol.%) (**b**), and POPC/SM/CHOL (60/20/20 mol.%) (**c**) mediated by 40 mM CaCl_2_, 20 wt.% PEG-8000, 150 μm FP_SARS(816-827)_, respectively. The relationship between the color, the number, and the CLP is given in the upper panel (**c**).

Next, we conducted a fusion inhibition assay with CLP-pretreated vesicles composed of DOPC/CHOL (80/20 mol.%) and PEG-8000 as a fusion trigger (Figure 2b, upper panel). Only three CLPs, aculeacin A, caspofungin, and fengycin, were able to suppress the fusion of DOPC/CHOL liposomes mediated by PEG-8000, whereas the other CLPs did not significantly affect the maximum value of *% fusion*. (Supplementary Table S2).

Additionally, we examined the effects of CLPs on the fusion of vesicles composed of POPC/SM/CHOL (60/20/20 mol.%) mediated by a short fragment of SARS-CoV-2 fusion peptide 1 (816-SFIEDLLFNKVT-827, FP_SARS(816-827)_) (Figure 2c, upper panel). Similar to the calcium-mediated fusion of DOPC/DOPG/CHOL liposomes, aculeacin A, anidulafungin, caspofungin, fengycin, iturin A, mycosubtilin, syringostatin A, and syringotoxin B were shown to reduce the *% fusion* of POPC/SM/CHOL (60/20/20 mol.%) vesicles induced by FP_SARS(816-827)_. The efficiency of CLPs increased in the series anidulafungin ≈ aculeacin A < iturin A ≤ syringostatin A ≈ syringotoxin B ≤ caspofungin ≈ fengycin (Supplementary Table S2). Interestingly, in contrast to the results obtained in previous systems, daptomycin showed slight inhibition, while polymixin B notably induced the FP_SARS(816-827)_-mediated fusion of SM-enriched liposomes (Supplementary Table S2). We cannot fully exclude the possibility that CLPs could interact with the SARS-CoV-2 fusion peptide and prevent specific rearrangements of the lipid matrix under its action. However, docking analysis revealed that daptomycin and surfactin were characterized by approximately 2-fold better efficacy of binding with the spike glycoprotein (S protein) of SARS-CoV-2 than iturin A but 2-fold lower efficacy than the leading compounds [19]. These data are not consistent with the results presented in Supplementary Table S2, demonstrating that the revealed ability of CLPs to inhibit vesicle fusion under the action of FP_SARS(816-827)_ was not likely to be due to their specific binding to the fusion peptide.

Considering the results presented on upper panel of Figure 2 we concluded that the ability of CLPs to inhibit fusion depended strictly on the membrane composition. Taking into account the inverse relationship between *% fusion* and the efficiency of the antifusogenic action of CLPs, we chose another parameter for further consideration, *% inhibition*. We accepted that CLP should be considered an effective fusion inhibitor if the *% inhibition* was higher than 50%. Our data demonstrate that caspofungin and fengycin were the most versatile and effectively inhibited the fusion of DOPC/DOPG/CHOL, DOPC/CHOL, and POPC/SM/CHOL liposomes mediated by calcium, PEG-8000, and FP_SARS(816-827)_, respectively (Figure 2, lower panel). Iturin A, mycosubtilin, and syringotoxin B suppressed the fusion of both DOPC/DOPG/CHOL and POPC/SM/CHOL vesicles triggered by calcium and FP_SARS(816-827_), respectively (Figure 2a, b, lower panel). Aculeacin A was able to inhibit the fusion of both DOPC/DOPG/CHOL and DOPC/CHOL liposomes mediated by calcium and PEG-8000, respectively (Figure 2a, b, lower panel). Anidulafungin, syringostatin A and daptomycin were the most specific. Anidulafungin exclusively demonstrated antifusogenic activity in relation to the calcium-induced fusion of DOPC/DOPG/CHOL vesicles (Figure 2a, lower panel), while syringostatin A and daptomycin were able to suppress the FP_SARS(816-827)_-mediated fusion of POPC/SM/CHOL liposomes (Figure 2c, lower panel).

### Antiviral activity of CLPs against SARS-CoV-2

Furthermore, we examined whether the tested CLPs were active against SARS-CoV-2. Daptomycin, polymyxin B, and surfactin were toxic to *Vero* cells at all concentrations used and therefore were excluded from further study. Aculeacin A, anidulafungin, caspofungin, iturin A, and mycosubtilin were toxic at concentrations greater than 12.5, 6.25, 100, 50, and 12.5 μg/ml, respectively (Table 1).

**Table 1.** Cytotoxicity of the tested lipopeptides against *Vero* cells and their antiviral activity against SARS-CoV-2

| *CLP* | $C^{\&}$, $\mu\text{g/ml}$ | $\text{lgTCID}_{50}/\text{ml}^{\$}$ |
| --- | --- | --- |
| *Aculeacin A* | 12.5 | 3 |
| *Anidulafungin* | 6.25 | 2 |
| *Caspofungin* | 100 | 6 |
| *Iturin A* | 50 | 3 |
| *Mycosubtilin* | 12.5 | 3 |
| *Surfactin* | toxic in all tested concentrations | – |
| *Daptomycin* |  |  |
| *Polymyxin B* |  |  |

& – The maximum non-toxic concentration of CLPs
§ – The $\text{lgTCID}_{50}/\text{ml}$ in the absence of CLPs was equal to 6.

To determine the infectious activity of viral progeny, we prepared tenfold dilutions of the supernatants collected from the experimental wells incubated with CLPs and viral controls in serum-free DMEM/F12 medium. The resulting dilutions were added to a 96-well culture plate with an 80-90% *Vero* cell monolayer and incubated for 72 hours. After that, visual assessment of the virus-specific cytopathogenic effect was performed using a CKX41 inverted microscope. The virus titer in control specimens without CLPs was 10^6^ TCID_50_/ml. Caspofungin did not demonstrate antiviral activity against SARS-CoV-2. Aculeacin A, iturin A, and mycosubtilin at the maximum nontoxic concentrations were able to decrease the viral titer by 3 orders of magnitude (Table 1). The infectivity of the supernatant of cells treated with 6.25 μg/ml anidulafungin was 10^4^-fold lower than that of untreated cells (Table 1). The antiviral activity of aculeacin A, anidulafungin, iturin A, and mycosubtilin was qualitatively consistent with their ability to suppress the fusion of POPC/SM/CHOL vesicles induced by FP_SARS(816-827)_ (Figure 2c). At the same time, quantitative disagreements were observed in the effectiveness of the antiviral and inhibitory action of CLPs, which might be related to a variety of mechanisms of the antiviral action of CLPs. The antiviral activity of anidulafungin against SARS-CoV-2 was also shown in [20]. Using molecular dynamics methods, Ahamad et al. [21] suggested that anidulafungin targeted ACE2-spike protein interactions. Based on the *in silico* results, Dey et al. [22] associated the anti-SARS-CoV-2 action of anidulafungin with its interaction with the nsp12 protein, which controls the RNA-dependent RNA polymerase activity of SARS-CoV-2. Using molecular docking methods, Anwaar et al. [23] demonstrated the possibility of binding anidulafungin to a number of SARS-CoV-2 viral proteins: RNA-dependent RNA polymerase (RdRp), helicase (Hel), guanine-N7 methyltransferase (ExoN), uridylate-specific endoribonuclease, and surface glycoproteins S and N.

### Mechanisms of the antifusogenic action of CLPs

#### CLP-induced alteration in membrane electrostatic properties

At neutral pH, caspofungin, fengycin, syringostatin A, syringotoxin B, and polymyxin B have a net positive electric charge, while surfactin and daptomycin are negatively charged (Figure 1). Considering the key role of electrostatic interactions in the initial stages of the fusion of negatively charged liposomes under the action of calcium cations, it can be assumed that positively charged CLPs should slow down the calcium-mediated fusion of DOPG-enriched vesicles, while negatively charged CLPs should act in the opposite manner. To examine this idea, we analyzed the kinetics of vesicle fusion (Supplementary Table S3). The Pearson coefficient of correlation between the time constant, *τ*, characterizing the exponential dependence of calcein leakage at calcium-mediated fusion of DOPC/DOPG/CHOL liposomes on time and the net charges of the tested CLP molecules, was equal to 0.5. This confirmed the importance of charge–charge interactions in the system, although the weak correlation indicated that this was not the only factor affecting the characteristic time.

The short fragment of SARS-CoV-2 fusion peptide 1 used, FP_SARS(816-827)_, has a net negative charge. This means that the preincubation of electrically neutral liposomes composed of POPC/SM/CHOL with positively and negatively charged CLPs should promote and arrest the peptide interaction with the membrane, respectively. No correlation was observed between the time constant, *τ*, characterizing the exponential dependence of calcein leakage at FP_SARS(816-827)_-mediated fusion of POPC/SM/CHOL vesicles on time and the net charges of the tested CLP molecules, indicating that charge–charge interactions were not of key importance in alteration of peptide-induced fusion by CLPs. However, this did not exclude the probability of influence of CLPs on the electrostatic interactions between the fusion peptide and the membrane via dipole potential modulation. For example, the alteration in the membrane dipole potential was shown to affect the extent of the membrane fusion caused by the simian immunodeficiency virus fusion peptide and implicated the dipolar properties of membranes in the fusion [24]. Moreover, the S2 SARS-CoV fusion protein was shown to reduce the dipole potential of the bilayer [25]. To test the possibility we additionally examined the effects of CLPs on the boundary potential of POPC/SM/CHOL (60/20/20 mol.%) bilayers. Supplementary Table S4 shows that the changes in the membrane boundary potential upon the addition of anidulafungin, caspofungin, fengycin, iturin A, surfactin, syringostatin A, syringotoxin B, daptomycin, and polymyxin B at the concentrations used in a fusion assay did not exceed 16 mV. This indicates the inability of these CLPs to alter the boundary potential of POPC/SM/CHOL bilayers (and its components, including dipole potential). The addition of aculeacin A up to 1 μM significantly reduced the membrane boundary potential by 70 mV (Supplementary Table S4). There was no correlation between the *% inhibition* of POPC/SM/CHOL vesicles in the presence of CLPs (Figure 2c, lower panel) and CLP-induced changes in the boundary potential of POPC/SM/CHOL membranes (Supplementary Table S4). In summary, we can conclude that the CLPs did not interfere with membrane fusion via their electrostatic effects.

#### CLP-induced alteration in lipid packing

Figure 1 shows that the CLPs tested differ not only in the charge of their peptide heads but also in the length of the hydrocarbon tails. We noticed that as a rule, the longer the tail, the higher the antifusogenic activity (Figure 3a, the corresponding Pearson coefficient of correlation between the *% inhibition* of calcium-mediated fusion of DOPC/DOPG/CHOL liposomes and the number of carbon atoms bound into a linear chain is equal to 0.7) and suggested that CLPs affect membrane fusion by altering lipid packing. A similar conclusion was drawn in our recent study of the effects of alkaloids on the fusion of lipid vesicles [1].

**Figure 3.**
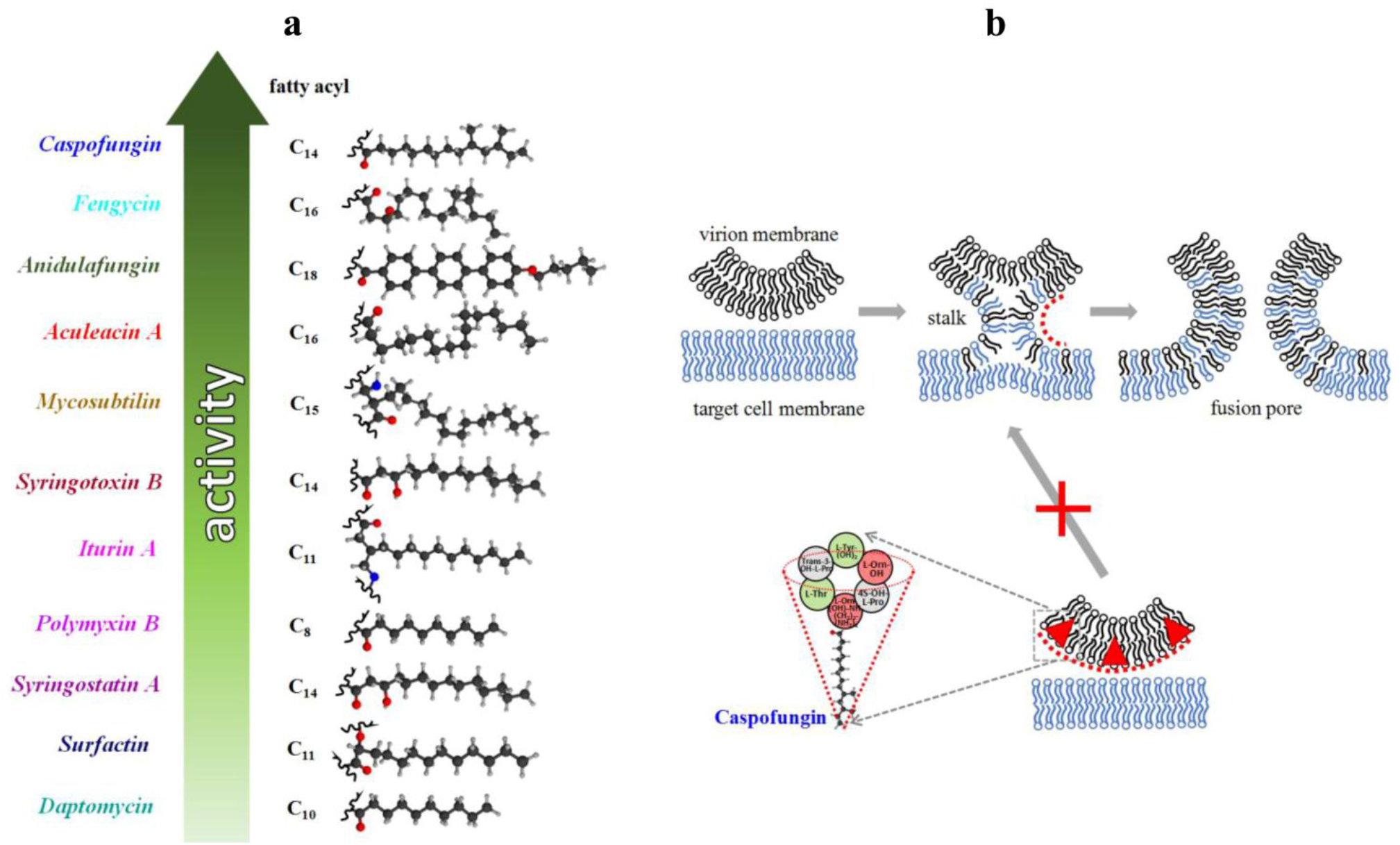
(**a**) Schematic representation of the relationships between the *% inhibition* of calcium-mediated fusion of DOPC/DOPG/CHOL (40/40/20 mol.%) liposomes and the length of the hydrocarbon tail of the CLPs. (**b**) Schematic representation of membrane fusion between viral and target cell membranes: the formation of an intermediate state, called the “stalk”, and a fusion pore are proposed. The addition of CLP molecules (red triangles) led to the inhibition of membrane fusion due to the higher energetic costs of the formation of lipid surfaces of negative curvature.

To gain mechanistic insight into the CLP-induced changes in lipid ordering, we conducted DSmC measurements with a DPPC/CHOL (90/10 mol.%) mixture. The melting temperature (*T_m_*) of the untreated vesicles was equal to 40.6 ± 0.1°C. The width of the melting peak at half-height (*T_1/2_*), which is related to sharpness of the gel-to-liquid-crystalline (L_β_/L_α_) phase transition, was approximately 0.7°C. Figure 4 shows that aculeacin A (a), anidulafungin (b), caspofungin (c), fengycin (d), and iturin A (e) were able to affect DPPC/CHOL melting in a dose-dependent manner, whereas the surfactin (f), daptomycin (g), and polymyxin B (h) had negligible ability to modulate DPPC/CHOL thermotropic behavior. The ability of CLPs to reduce *T_m_* increased in the following series: aculeacin A ≤ caspofungin ≤ iturin A < anidulafungin << fengycin at all tested lipid:CLP ratios (50:1, 25:1, and 10:1) (Table 2). The ability to enhance *T_1/2_* increased in the following series: iturin A ≈ aculeacin A ≤ anidulafungin ≈ caspofungin < fengycin (Table 2). In addition to increasing *T_1/2_*, the decrease in melting cooperativity was expressed as a 1.3-1.5-fold drop in the cooperative unit size (*CUS*) in the presence of caspofungin at lipid:CLP ratios of 25:1 and 10:1 and fengycin at lipid:CLP ratios of 50:1 and 25:1 (Table 2). Good profile matching from two repeated heating steps (Δ*T_m1-m2_* was close to zero, Table 2) indicated the equilibrium distribution of CLPs between lipid monolayers. The introduction of anidulafungin, caspofungin, and fengycin up to low lipid:CLP ratios (25:1 and 10:1) turned the main peak into a multicomponent profile consisting of two overlapping transitions (Figure 4b,c,d, insets). This might indicate the formation of at least two lipid phases with different concentrations of CLPs: the right component (peak #1), characterized by a higher melting point, might be attributed to a practically pure DPPC/CHOL mixture, and the left component (peak #2), characterized by a lower melting point, should be related to CLP-enriched lipid domains. The invariability in the *T_m_*-hysteresis (Δ*T_h_*, the difference in the melting temperature between heating and cooling scans) with increasing concentrations of CLPs indicated the absence of lipid interdigitation in the presence of CLPs (Table 2).

**Figure 4.**
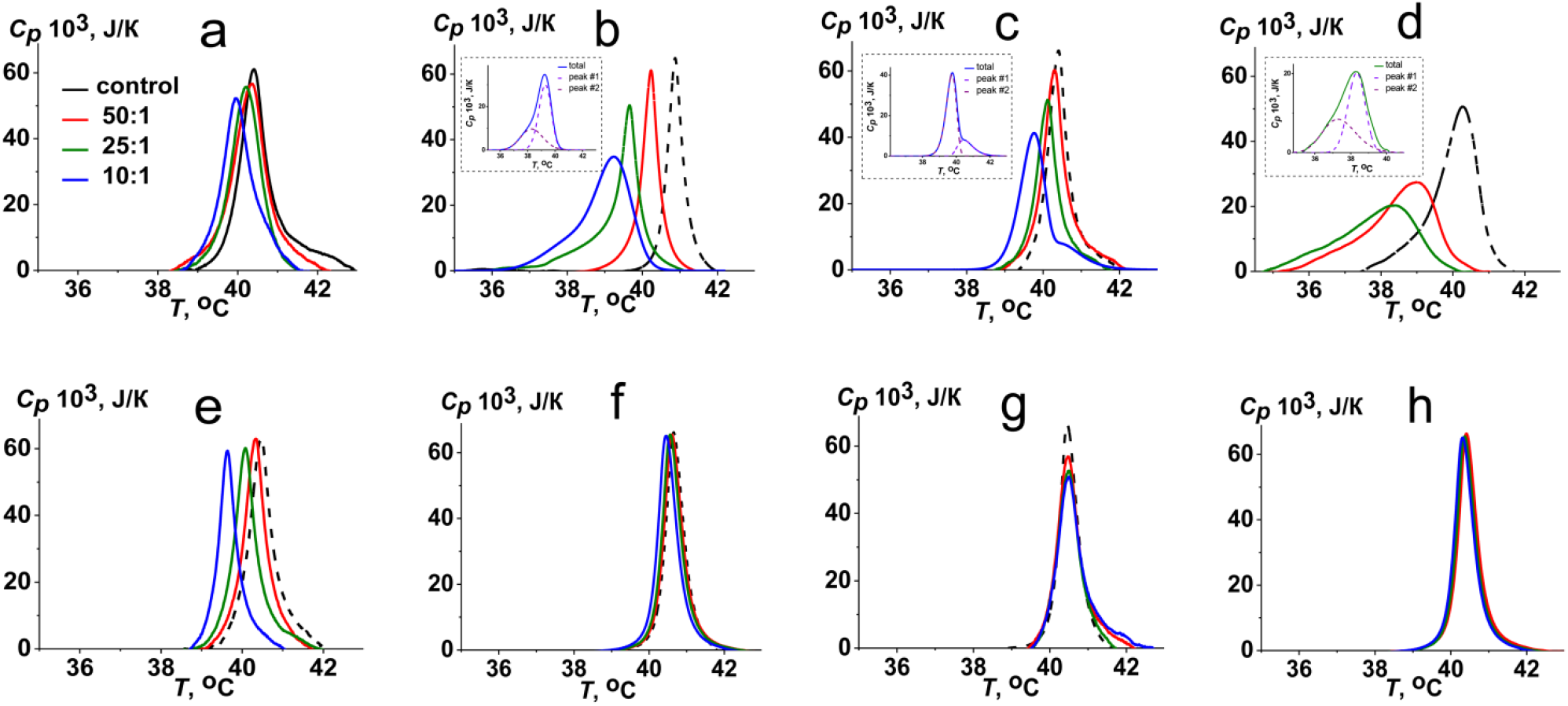
Heating thermograms of the gel-to-liquid-crystalline (L_β_/L_α_) phase transition of DPPC/CHOL (90/10 mol.%) in the absence (control, *black curves*) and presence of aculeacin A (**a**), anidulafungin (**b**), caspofungin (**c**), fengycin (**d**), iturin A (**e**), surfactin (**f**), daptomycin (**g**), and polymyxin B (**h**). The lipid:CLP molar ratio is equal to 50:1 (*red curves*), 25:1 (*green curves*) and 10:1 (*blue curves*). *Insets*: Deconvolution analysis of the L_β_/L_α_ transition peak of DPPC/CHOL (90/10 mol.%) in the presence of anidulafungin (**b**), caspofungin (**c**), and fengycin (**d**) at lipid:CLP molar ratios of 10:1, 10:1, and 25:1, respectively.

**Table 2.**
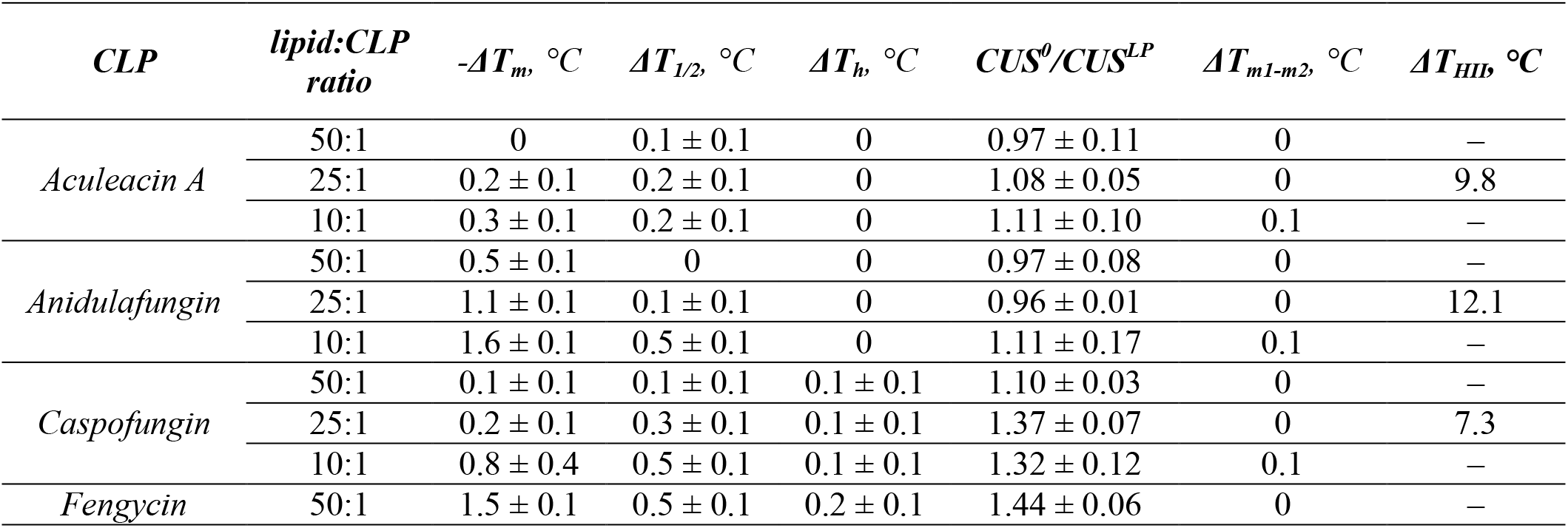
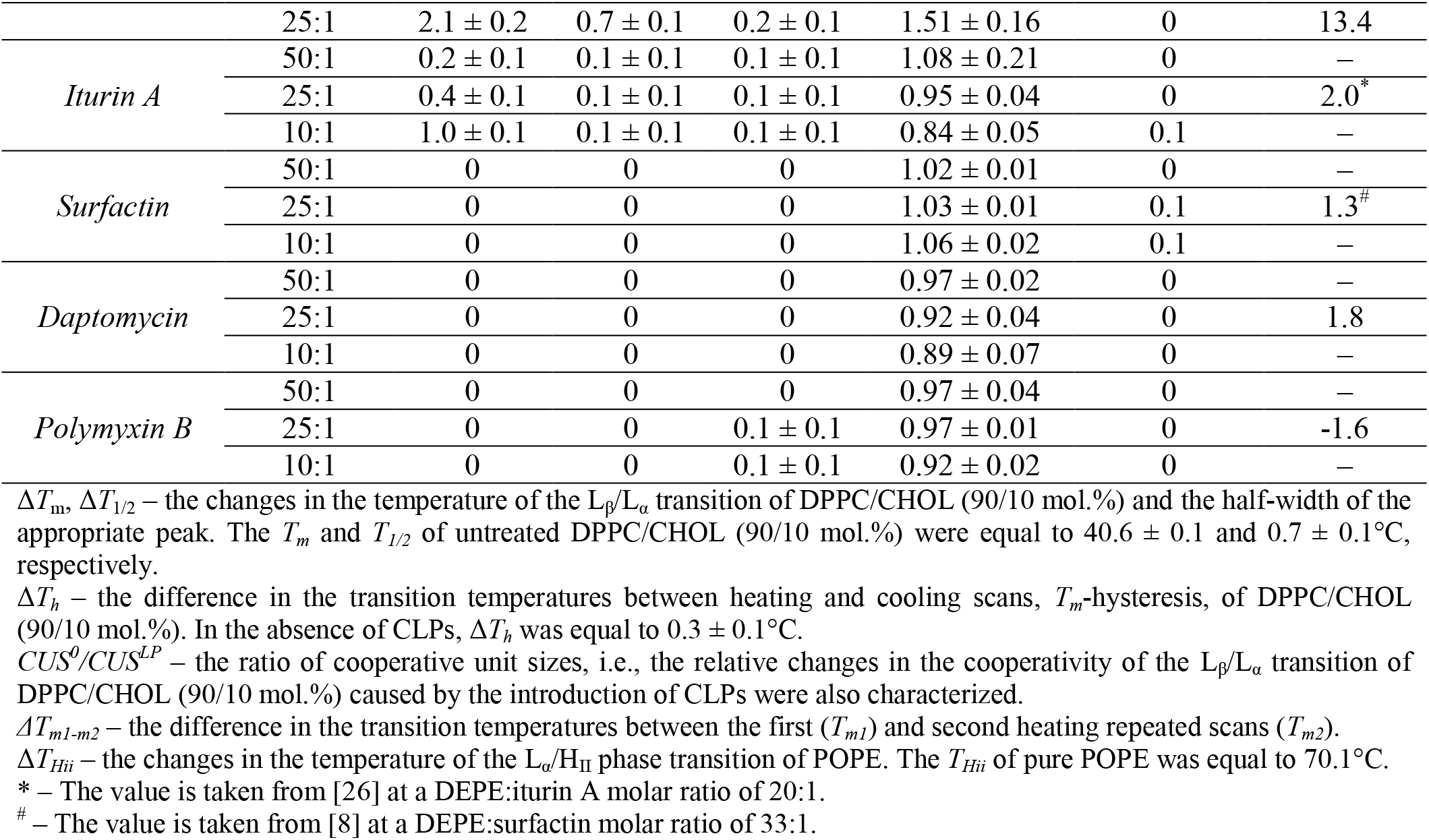
The effect of CLPs on the thermotropic behavior of membrane-forming lipids

The Pearson correlation coefficient between the *% inhibition* of the CLPs of CaCl_2_-, PEG-8000- and FP-SARS-CoV-2-induced liposome fusion and the Δ*T_1/2_*-values at all tested lipid:CLP ratios varied in the range of 0.5-0.9, indicating the possible relation of antifusogenic and disordering abilities of CLPs.

Another possible mechanism for the antifusogenic action of CLPs might be related to their ability to induce positive curvature stress in lipid bilayers due to the conical shape of CLP molecules [8,27]. The induction of positive curvature stress by CLPs might increase the energetic costs of the formation of negatively curved structures, especially fusion intermediates such as stalks (Figure 3b). To test this possibility, we examined the lamellar-to-inverted hexagonal (L_α_/H_II_) phase transition of POPE using differential scanning microcalorimetry (Figure 5a, Table 2). In the absence of CLPs, the low-enthalpy L_α_/H_II_ transition of POPE occurred at approximately 70°C (Figure 5a, control). Caspofungin, aculeacin A, anidulafungin, and fengycin at a lipid:CLP molar ratio of 25:1 increased the transition temperature by 7.3, 9.8, 12.1, and 13.4°C, respectively (Figure 5a, Table 2). Iturin A and surfactin at molar ratios of 20:1 and 33:1 are known to increase the transition temperature of DEPE by 2.0 and 1.3°C, respectively [8,26] (Table 2). Moreover, daptomycin and polymyxin B at a lipid:CLP molar ratio of 25:1 were found to slightly increase and diminish the *T_Hii_* of POPE by 1.8 and 1.6°C, respectively (Table 2). A significant increase in the L_α_/H_II_ transition temperature in the presence of caspofungin, aculeacin A, anidulafungin, and fengycin indicated that the CLPs suppressed the formation of lipid structures with high negative curvature, and this ability of CLPs correlated with their inhibitory effect on membrane fusion triggered by CaCl_2_, PEG-8000, and FP_SARS(816-827)_ (Pearson’s correlation coefficients were equal to 0.8, 0.7, and 0.5, respectively). Figure 5b shows a schematic representation of the inhibition of the L_α_/H_II_ transition of membrane-forming lipids by CLP.

**Figure 5.**
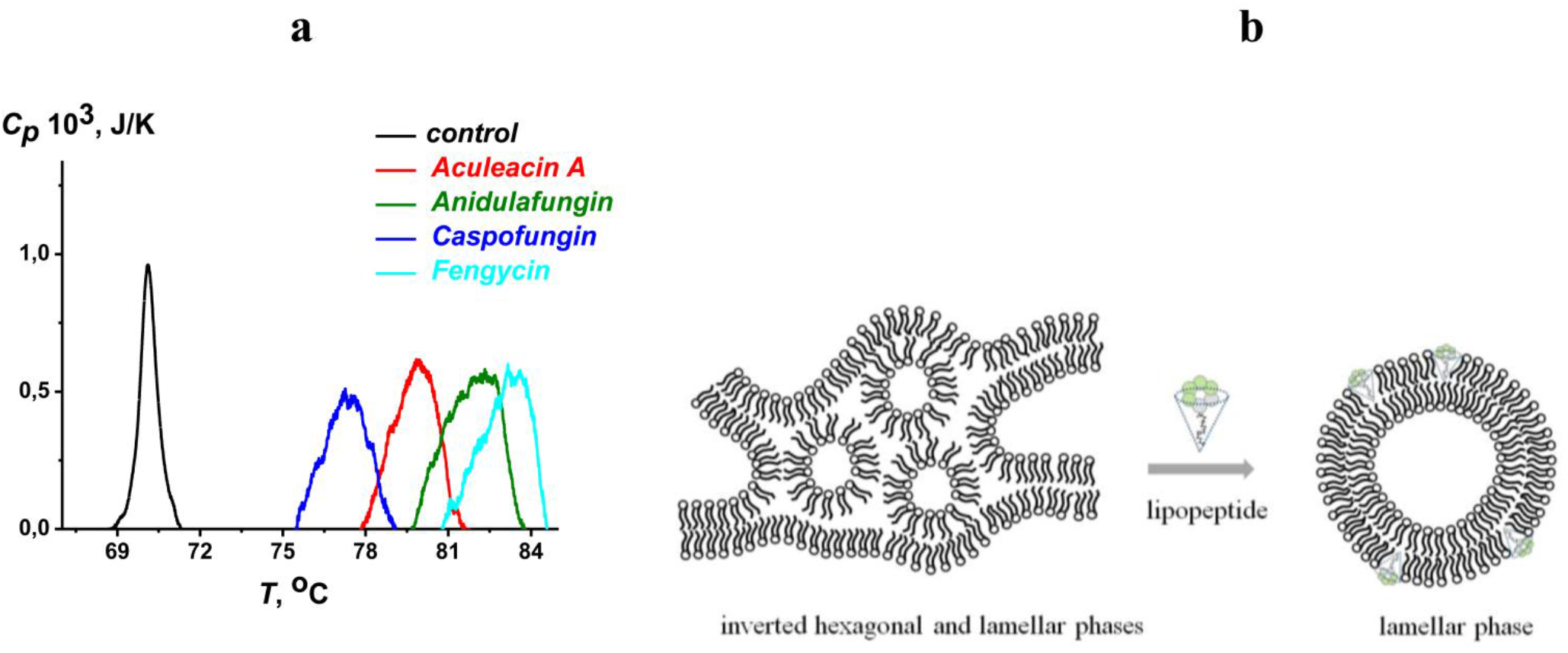
(**a**) Thermograms of the lamellar-to-inverted hexagonal (L_α_/H_II_) phase transition of POPE in the absence (control, *black curve*) and presence of aculeacin A (*red curve*), anidulafungin (*olive curve*), caspofungin (*blue curve*), and fengycin (*cyan curve*) at a lipid:CLP ratio of 25:1. (**b**) Schematic representation of the transition from the inverted hexagonal phase to the lamellar phase of POPE in the presence of aculeacin A, anidulafungin, caspofungin, and fengycin.

The correlation coefficient between Δ*T_Hii_*-values and *% inhibition* of FP_SARS(816-827)_-induced fusion of POPC/SM/CHOL (60/20/20 mol.%) vesicles was lower than the correlation coefficient between Δ*T_Hii_*-values and *% inhibition* of CaCl_2_ and PEG-8000-mediated fusion of DOPC/DOPG/CHOL (40/40/20 mol.%) and DOPC/CHOL (80/20 mol.%) liposomes might be explained if the first type of membranes, under appropriate conditions, can exhibit lipid phase separation and undergo lipid raft formation [28], while the last two do not.

There is much evidence that lipid rafts might be involved in fusion with the target cell membrane of a variety of enveloped viruses, including HIV-1 [29], HSV-1 [30], Ebola and Marburg [31,32], influenza [33], and Newcastle disease virus [34]. In the cases of HIV-1, HSV-1, and Newcastle disease virus, the major role in the fusogenic activity of viral peptides was attributed to cholesterol [29,30,34]. Recent data also indicate that CHOL-enriched membrane domains are of key importance for SARS-CoV-2 entry into cells [35–39]. Moreover, viral fusion peptides, particularly that of HIV, preferentially target the L_o_/L_α_ boundary regions [40,41].

Risselada [42] showed that the location of highly negatively curved fusion intermediates, stalks, near the L_o_/L_α_ phase boundary depends on the spontaneous curvature of L_o_ phase constituents: the more negative the curvature is, the higher the probability of stalk formation, and vice versa. The L_o_ domains (lipid rafts) are characterized by a larger hydrophobic thickness than the surrounding L_α_ regions [43–45]. The CLPs producing positive curvature might compensate for the hydrophobic mismatch and a negative curvature stress at a L_o_/L_α_ phase boundary, minimizing the free energy of raft formation and increasing the fusion energy barrier. This assumption is consistent with the preferential partitioning of amphipathic peptides to the boundary of L_o_ domains [46,47]. Numerous literature data on the modulation of the domain organization of model and cell membranes by CLPs are also consistent with this assumption. Thus, the preferential distribution of syringomycin E (a close analog of syringostatin A and syringotoxin B) into lipid rafts was shown [48]. Wójtowicz et al. [49] demonstrated that surfactin targets raft domains. Fengycin and polymyxin B were shown to modulate the phase separation and domain organization of microbial lipids [50,51]. Mycosubtilin induced changes in the organization and morphology of cholesterol- and sphingomyelin-enriched membranes [52]. Membrane reorganization by daptomycin is intensively discussed [53]. Müller et al. [54] supposed that daptomycin modifies L_α_ domains in the *Bacillus subtilis* membrane. Caspofungin-induced lipid and protein redistribution in the plasma membrane of *C. albicans* [55,56] also suggests that this CLP influences membrane lipid segregation and domain formation.

In line with this idea, we tested the possibility that the CLPs affect the phase separation in the SM-enriched membrane (Supplementary Figure S2). Figure 6a presents the effects of the tested CLPs at a lipid:CLP molar ratio of 25:1 on the area fraction of lipid rafts in giant vesicles composed of DOPC/SM/CHOL (40/40/20 mol.%) (S_0_). It should be noted that FP_SARS(816-827)_ at a lipid:peptide molar ratio up to 10:1 did not affect the macroscopic domain organization of the liposomes (data not shown). Anidulafungin, caspofungin, and fengycin were shown to significantly increase S_0_. Data from DSmC showed that some CLPs, especially anidulafungin, caspofungin, and fengycin, were able to disorder membrane lipids (decrease *T_m_* and increase *T_1/2_*) (Figure 4, Table 2). Taking into account the disordering action of CLPs, three different ways to change the domain organization of membranes can be proposed: (i) to fluidize the L_α_ domains, to increase the hydrophobic mismatch at the L_o_/L_α_ phase boundary, and to diminish the probability of raft formation (Figure 6b I); (ii) to disorder ordered domains, to reduce the hydrophobic mismatch at the L_o_/L_α_ phase boundary, and to enhance raft formation (Figure 6b II) (the DSmC data obtained with saturated high melting DPPC/CHOL were consistent with this possibility); and (iii) to induce positive curvature at the L_o_/L_α_ phase boundary and to increase the probability of raft formation (Figure 6b III) (the DSmC data obtained with nonlamellar POPE were consistent with this possibility).

**Figure 6.**
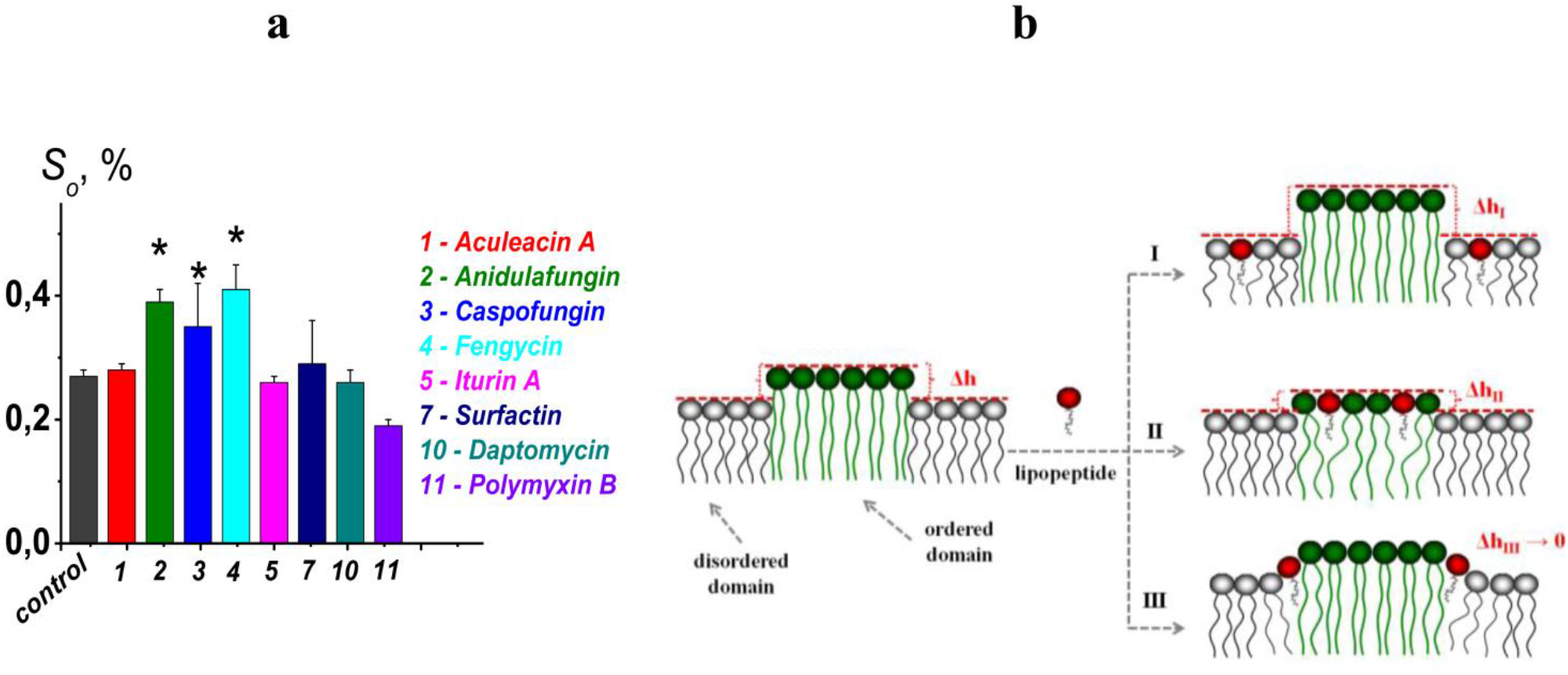
(**a**) The area fraction of lipid rafts (S_0_, %) in giant vesicles composed of DOPC/SM/CHOL (40/40/20 mol.%) with a lipid:CLP molar ratio of 25:1. * – *p* ≤ 0.01 (Mann–Whitney-Wilcoxon’s test, control *vs*. CLP) (**b**) The schematic representation of the CLP action on L_o_/L_α_ phase separation in the bilayer: (I) the disordering action of CLP on L_α_ domains and increasing hydrophobic mismatch at the L_o_/L_α_ phase boundary (Δh_I_ > (Δh), (II) the disordering action of CLP on L_o_ domains and decreasing hydrophobic mismatch at the L_o_/L_α_ phase boundary (Δh_II_ < Δh), (III) inducing positive curvature stress by CLP at L_o_/L_α_ phase boundary.

We cannot discriminate between the last two possibilities because of the potentiation of lipid raft formation by anidulafungin, caspofungin, and fengycin (Figure 6a), which have the greatest effects on both DPPC/CHOL melting (Figure 4b,c,d, Table 2) and the lamellar-to-inverted hexagonal phase transition of POPE (Figure 5a, Table 2). Moreover, these CLPs demonstrated larger inhibitory activity against the membrane fusion of liposomes composed of DOPC/DOPG/CHOL (40/40/20 mol.%) mediated by calcium (Figure 2, lower panel, a), while caspofungin and fengycin were also effective against the PEG-8000- and FP_SARS(816-827)_-induced fusion of vesicles made from DOPC/CHOL (80/20 mol.%) and POPC/SM/CHOL (60/20/20 mol. %) (Figure 2b,c, lower panel). Although a precise understanding of the mechanism by which CLPs control virus infectivity requires future structural analysis a strong interrelation between the abilities of CLPs to produce positive curvature stress, inhibit membrane fusion and disorder membrane lipids should be underlined. More probable, CLPs are deeply embedded into bilayer by incorporating their relatively long tails into hydrocarbon core, while large hydrophilic heads increase the lateral pressure in the lipid head group region. This, in turn, causes both the production of positive curvature stress and an increase in the energy of formation of structures with reverse curvature (inverted hexagonal phases and stalks), and also leads to an increase in the area per lipid molecule and a subsequent increase in the conformational mobility of lipid hydrocarbon chains.

In conclusion, our study provides insights into the critical role in membrane fusion of alterations in the physical properties of the lipid environment and fusion peptide-lipid interactions. Lipid-mediated inhibition of the fusion of the virus lipid envelope with cell membranes holds promise for developing alternative antiviral strategies. Aculeacin A, anidulafugin, iturin A, and mycosubtilin are proposed to prevent and combat SARS-CoV-2 infection, and experimental validation in preclinical studies is needed. Our data pave the way for targeting the inhibitors not to the interaction of viral protein with the cell receptor, but to the fusion of the viral and cell membranes that can open serious advantages for reducing both an emergence of viral resistance and strain-specificity to drugs with lipid-associated mechanism of action.

## Materials and Methods

### Materials

Calcein, nonactin, HCl, KCl, NaCl, CaCl_2_, HEPES, KOH, NaOH, PBS, hexadecane, pentane, ethanol, methanol, Triton X-100, DMSO, Sephadex G-50, sorbitol, polyethylene glycol 8000 (PEG-8000), and cyclic lipopeptides (aculeacin A, anidulafungin, caspofungin, fengycin, iturin A, mycosubtilin, surfactin, daptomycin, and polymyxin B) were purchased from Sigma–Aldrich Company Ltd. (Gillingham, United Kingdom). Syringostatin A and syringotoxin B were kindly provided by Dr. D. Takemoto (Utah State University, USA).

The lipids 1-palmitoyl-2-oleoyl-*sn*-glycero-3-phosphocholine (POPC), 1,2-dipalmitoyl-*sn*-glycero-3-phosphocholine (DPPC), 1-palmitoyl-2-oleoyl-*sn*-glycero-3-phosphoethanolamine (POPE), 1,2-dioleoyl-*sn*-glycero-3-phosphocholine (DOPC), 1,2-dioleoyl-*sn*-glycero-3-phosphoglycerol (DOPG), cholesterol (CHOL), sphingomyelin (brain, porcine) (SM), and 1,2-dipalmitoyl-*sn*-glycero-3-phosphoethanolamine-N-(lissamine rhodamine B sulfonyl) (Rh-DPPE) were obtained from Avanti Polar Lipids^®^(Avanti Polar Lipids, Inc., USA).

A short fragment of SARS-CoV-2 fusion peptide 1 (816-SFIEDLLFNKVT-827, FP_SARS(816-827)_) was commercially synthesized (IQ Chemical, Russia). The purity of the peptide was ≥ 98%. The product was purified by column chromatography, and the final compound was fully characterized by mass spectrometry. Taking into account the key role of the LLF motif in membrane rearrangements during peptide-induced fusion [57,58], a peptide composed of SFIEDAAANKVT was used as a negative control.

*Vero* monkey kidney epithelial cells (ATCC CCL81) were used to analyze the antiviral activity of the compounds. DMEM, DMEM/F12 culture medium, fetal bovine serum (FBS), trypsin solution, and Dulbecco’s phosphate-buffered saline (DPBS) were manufactured by Gibco (Gibco^™^, Life Technologies, Scotland).

### Membrane Fusion Assay

Large unilamellar vesicles (Ø100 nm) composed of DOPC/CHOL (80/20 mol.%), DOPC/DOPG/CHOL (40/40/20 mol.%) and POPC/SM/CHOL (60/20/20 mol.%) were obtained by extrusion using an Avanti Polar Lipids^®^ mini-extruder (Avanti Polar Lipids, Inc., USA). The lipid stock solution in chloroform was dried under a gentle stream of nitrogen. The dry lipid film was hydrated by a buffer (35 mM calcein, 10 mM HEPES, pH 7.4). The suspension was subjected to five freeze–thaw cycles and then passed through a 100 nm Nuclepore polycarbonate membrane 13 times. The liposome suspension contained 3 mM lipid. The calcein that was not entrapped in the vesicles was removed by gel filtration with a Sephadex G-50 column to replace the buffer outside the liposomes with calcein-free solution (150 mM NaCl, 10 mM HEPES, pH 7.4). Calcein in vesicles fluoresces very poorly because of its strong self-quenching at millimolar concentrations. Dye leakage upon vesicle fusion is designed to indicate the level of mixing of the internal content of liposomes and is used to quantify vesicle fusion [59]. Here, 40 mM CaCl_2_, 20 wt.% PEG-8000, and 150 μM FP_SARS(816-827)_ were used to induce liposome fusion. To test the fusion inhibitory activity of CLPs, CaCl_2_, PEG-8000 or FP-SARS-CoV-2 was added to a suspension of vesicles preincubated with CLPs for 35 ± 5 minutes at the concentrations indicated in Supplementary Table S1. CLPs alone produced minimal calcein release from liposomes of appropriate composition at the indicated concentrations (the leakage did not exceed 10%). The relative intensity of fluorescence of calcein leaked upon vesicle fusion (*RF*, %) was determined using a Fluorat-02-Panorama spectrofluorimeter (Lumex, Saint-Petersburg, Russia). The excitation and emission wavelengths were 490 and 520 nm, respectively. All experiments were performed at room temperature (25°C).

The percentage of membrane fusion (*% fusion*) was calculated using the following equation:

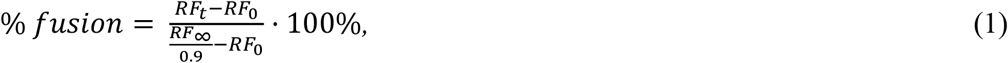

where *RF*_0_, *RF*_t_, and *RF*_∞_ are the fluorescence intensities at time = 0, time = *t* and time = ∞, respectively. *RF*_∞_ was measured in the presence of Triton X-100, which induced 100% disengagement of the marker by disrupting liposome membranes. A coefficient of 0.9 was introduced to take into account the dilution of the sample by detergent.

The dependence of *% fusion* on time was fitted using a single exponential function with time constant τ.

The antifusogenic activity of CLPs was presented as the extent of fusion inhibition (*% inhibition*). The parameters were averaged from 2 to 5 independent experiments and presented as the *mean* ± *standard error* (*p* ≤ 0.01).

### Antiviral Testing

To analyze the antiviral activity of the compounds, *Vero* monkey kidney epithelial cells (ATCC CCL81) were grown in culture flasks (Nunc, Denmark) in DMEM supplemented with 10% FBS. *Vero* cells were detached from the flask surface using 0.25% trypsin solution. The cell concentration was adjusted to 5·10^5^ cells/ml, and cells were seeded into 96-well culture plates (0.1 ml per well). The plates were incubated for 24 h at 37°C in an atmosphere of 5% CO_2_. To assess the antiviral activity of CLPs, SARS-CoV-2 (isolate 17612) with an infectious titer of 10^3^ TCID_50_/ml was used. CLPs were dissolved in DMSO, and then twofold serial dilutions from 200 to 1.56 μg/ml in serum-free DMEM/F12 were prepared. Afterward, 100 μl/well of each dilution of CLP was added to *Vero* cells in the appropriate wells of culture plates and incubated for 1 h at 37°C in a 5% CO_2_ atmosphere. Serum-free DMEM/F12 was added to the control wells instead of CLPs. After 1 h of incubation, 100 μl/well SARS-CoV-2 was added to the wells and incubated for 1 h at 37°C in a 5% CO_2_ atmosphere. The virus-containing fluid was added to the wells of the viral control, and serum-free DMEM/F12 was added to the cell control. After incubation, the virus and compounds were removed, the cells were washed three times with DPBS, and 100 μl/well of each 200-1.56 μg/ml dilution of CLPs was added to the corresponding wells. Serum-free DMEM/F12 was added to the control wells, and the plate was incubated for 24 h at 37°C in a 5% CO_2_ atmosphere. After 24 h, supernatant from the experimental wells and control wells with virus was collected and titrated by TCID_50_. For titration, *Vero* cells were used. The results of titration were evaluated after 72 h to determine the cytopathogenic effect of the virus. The viral titer was considered the highest dilution of viral harvest causing the appearance of virus-specific cytopathogenic effects.

### Membrane Boundary Potential Measurements

Virtually solvent-free planar lipid bilayers were prepared using a monolayer-opposition technique [60] on a 50-μm diameter aperture in a 10-μm thick Teflon film separating the two (*cis*- and *trans*-) compartments of the Teflon chamber. The aperture was pretreated with hexadecane. Lipid bilayers were made from POPC/SM/CHOL (60/20/20 mol.%). The steady-state conductance of K^+^-nonactin was modulated via the two-sided addition of CLPs at the concentrations indicated in Supplementary Table S1 (third column).

Ag/AgCl electrodes with 1.5% agarose/2 M KCl bridges were used to apply voltage (*V*) and measure the transmembrane current. Here, “positive voltage” refers to the case in which the *cis*-side compartment is positive with respect to the *trans*-side. The current was measured using an Axopatch 200B amplifier (Molecular Devices, LLC, Orleans Drive, Sunnyvale, CA, USA) in voltage clamp mode. Data were digitized using a Digidata 1440A and analyzed using pClamp 10.0 (Molecular Devices, LLC, Orleans Drive, Sunnyvale, CA, USA) and Origin 8.0 (OriginLab Corporation, Northampton, MA, USA). Data were acquired at a sampling frequency of 5 kHz using low-pass filtering at 1 kHz. All experiments were performed at room temperature (25°C).

The conductance of the lipid bilayer was determined by measuring the transmembrane current at a constant transmembrane voltage (*V* = 50 mV). The subsequent calculations of CLP-induced changes in the membrane boundary potential (Δ*φ_b_*) were performed according to the Boltzmann distribution [61].

The Δ*φ_b_*-values were averaged from 3 to 5 independent experiments and are presented as the *mean* ± *standard error* (*p* ≤ 0.05).

### Differential Scanning Microcalorimetry (DSmC)

#### 1) Gel-to-liquid crystalline phase transition

DSmC experiments were performed by a μDSC 7EVO microcalorimeter (Setaram, Caluire-et-Cuire, France). Giant unilamellar vesicles composed of DPPC/CHOL (90/10 mol.%) were prepared by the electroformation method using Vesicle Pre Pro^@^ (Nanion Technologies, Munich, Germany) (standard protocol, 3 V, 10 Hz, 1 h, 55°C). The resulting liposome suspension contained 2.5 mM lipid and was buffered by 5 mM HEPES at pH 7.4. The liposomal suspension was heated and cooled at a constant rate of 0.2 and 0.3 C min^-1^, respectively. CLPs were added to the liposome suspension up to lipid:CLP ratios of 50:1, 25:1, and 10:1. The reversibility of the thermal transitions was assessed by reheating the sample immediately after the cooling step from the previous scan. The temperature dependence of the excess heat capacity was analyzed using Calisto Processing (Setaram, Caluire-et-Cuire, France).

The thermograms of DPPC/CHOL (90/10 mol.%) were characterized by the melting temperature (the temperature at which excess heat capacity reaches a maximum, *T_m_*), the enthalpy of the main phase transition (an area of the main peak, Δ*H_cal_*), and *T_m_*-hysteresis (the difference in the transition temperatures between heating and cooling scans, Δ*T_h_*). The sharpness of the gel-to-liquid-crystalline (L_β_/L_α_) phase transition was expressed as the width of the peak at half-height (Δ*T_1/2_*). The changes in the cooperativity of the transition caused by the introduction of CLPs were also characterized by the ratio of cooperative unit sizes (*CUS*^0^/*CUS*^CLP^) determined according to [62].

The values of Δ*T_m_*, ΔΔ*H_cal_*, ΔΔ*T_h_*, Δ*T_1/2_*, and *CUS*^0^/*CUS*^CLP^ were averaged from 2 to 3 independent experiments and presented as the *mean* ± *standard deviation* (*p* ≤ 0.05).

#### 2) Lamellar-to-inverted hexagonal phase transition

The lamellar-to-inverted hexagonal (L_α_/H_II_) phase transition was proven to involve some of the same intermediate structures as membrane fusion [63], and we used DSmC of POPE to evaluate the effects of CLPs on negatively curved lipid formation. The POPE suspension was prepared by dissolving 8-9 mg of dry POPE without or with 0.35-0.45 mg of CLP in warm buffer (5 mM HEPES at pH 7.4). After that, the suspension was shaken in a water bath at 40°C for 60 min and sonicated at 40 kHz for 1-3 minutes. The resulting POPE suspension was refrigerated at 8°C for at least 8 hours before DSmC measurements. The POPE thermograms were characterized by the temperature of the L_α_/H_II_ phase transition.

### Confocal Fluorescent Microscopy

Giant unilamellar vesicles were prepared from a mixture of POPC/SM/CHOL (60/20/20 mol.%) and DOPC/SM/CHOL (40/40/20 mol.%) and 1 mol.% fluorescent lipid probe Rh-DPPE by the electroformation method (standard protocol, 3 V, 10 Hz, 1 h, 35°C) using Nanion Vesicle Prep Pro^@^ (Nanion Technologies, Munich, Germany). The resulting liposome suspension contained 1.5 mM lipid in 0.5 M sorbitol solution. The control samples were not modified. The liposome suspensions with and without CLPs were allowed to equilibrate for 30 min at room temperature. Vesicles were imaged through an oil immersion objective (65×/1.4HCX PL) using an Olympus (Hamburg, Germany). A helium-neon laser with a wavelength of 561 nm was used to excite Rh-DPPE. The temperature (25 ± 1°C) during observation was controlled by air heating/cooling in a thermally insulated camera.

The ability of FP_SARS(816-827)_ to fuse unilamellar vesicles composed of POPC/SM/CHOL (60/20/20 mol.%) was analyzed by the appearance of multilamellar and multivesicular liposomes (Supplementary Figure S1).

It is believed that Rh-DPPE clearly favors the liquid disordered (L_α_) phase and is excluded from the ordered phases, liquid ordered (L_o_, raft-like) and gel (L_β_) domains [64]. POPC/SM/CHOL (60/20/20 mol.%) at 23°C exhibits liquid ordered and gel (L_o_ + L_β_) phase separation [28] and therefore cannot be used for studying lipid raft dynamics in the presence of CLPs by confocal fluorescence microscopy of the distribution of Rh-DPPE. The DOPC/SM/CHOL (40/40/20 mol.%) mixture displays the coexistence of liquid disordered and liquid ordered (L_α_ + L_o_) phases at 23°C [65–67]. Thus, the uncolored lipid domains in vesicles composed of DOPC/SM/CHOL (40/40/20 mol.%) might be attributed to the L_o_ phase, i.e., to lipid rafts, while the red-colored domains might be assigned to the L_α_ phase labeled with Rh-DPPE. To quantify the area fraction of lipid rafts in giant vesicles, the ratios between the total area of all uncolored lipid domains in a single liposome and its area were estimated, the distribution of the area fraction of L_o_ domains over a population of 50 ÷ 320 vesicles from the same batch was constructed, and the *mean* ± *standard error* (*p* ≤ 0.05) was presented in the absence and presence of FP_SARS(816-827)_ and CLPs at lipid:(lipo)peptide molar ratios of 10:1 and 25:1, respectively.

## Acknowledgements

This research was supported in part by RSF (project N<u>o</u> 22-15-00417).

## Author contributions

E.V.S. and P.D.Z. performed membrane fusion assay, boundary potential measurements, and confocal fluorescent microscopy experiments. S.S.E. performed DSmC experiments. E.V.S., P.D.Z. and S.S.E. analyzed data. A.A.M., V.V.Z. and A.V.S. designed and performed antiviral testing experiments. O.S.O. designed experiments, supervised the study and wrote manuscript.

## Additional information

### Supplementary information

The online version contains supplementary material available.

## Supplementary information

**Supplementary Table S1:** Concentration of the tested lipopeptides used for DOPC/DOPG/CHOL (40/40/20 mol.%), DOPC/CHOL (80/20 mol.%), POPC/SM/CHOL (60/20/20 mol.%) with fusion caused by 40 mM CaCl2, 20 wt.% PEG-8000- or 150 μM FP_SARS(816-827)_.

| *lipopeptide* | **C, μM** |  |  |
| --- | --- | --- | --- |
|  | *CaCl<sub>2</sub>* | *PEG-8000* | *FP<sub>SARS(816-827)</sub>* |
| *Aculeacin A* | 2 | 2.5 | 1 |
| *Anidulafungin* | 20 | 5 | 10 |
| *Caspofungin* | 33 | 33 | 33 |
| *Fengycin* | 10 | 10 | 2 |
| *Iturin A* | 20 | 20 | 50 |
| *Mycosubtilin* | 25 | 25 | 25 |
| *Surfactin* | 0.01 | 0.01 | 0.01 |
| *Syringostatin A* | 20 | 20 | 20 |
| *Syringotoxin B* | 20 | 20 | 20 |
| *Daptomycin* | 100 | 100 | 100 |
| *Polymyxin B* | 1.25 | 10 | 10 |

**Supplementary Table S2.** The maximum percentage of fusion (%) of vesicles composed of DOPC/DOPG/CHOL (40/40/20 mol.%), DOPC/CHOL (80/20 mol.%), POPC/SM/CHOL (60/20/20 mol.%) mediated by 40 mM CaCl_2_, 20 wt.% PEG-8000- or 150 μM FP_SARS(816-827)_.

| *lipopeptide* | *CaCl<sub>2</sub>* | *PEG-8000* | *FP<sub>SARS(816-827)</sub>* |
| --- | --- | --- | --- |
| – | 81 ± 4 | 74 ± 5 | 69 ± 8 |
| *Aculeacin A* | 10 ± 6* | 9 ± 3* | 46 ± 14* |
| *Anidulafungin* | 4 ± 4* | 67 ± 2 | 48 ± 2* |
| *Caspofungin* | 1 ± 1* | 16 ± 2* | 5 ± 2* |
| *Fengycin* | 1 ± 1* | 22 ± 10* | 4 ± 3* |
| *Iturin A* | 39 ± 14* | 71 ± 7 | 21 ± 10* |
| *Mycosubtilin* | 11 ± 5* | 51 ± 12 | 11 |
| *Surfactin* | 86 ± 5 | 59 ± 4 | 68 ± 19 |
| *Syringostatin A* | 75 ± 1* | 66 ± 10 | 11 ± 1* |
| *Syringotoxin B* | 30 ± 6* | 71 ± 7 | 8 ± 3* |
| *Daptomycin* | 91 ± 3 | 63 ± 5 | 40 ± 17 |
| *Polymyxin B* | 51 ± 4* | 61 ± 7 | 86 ± 4* |

\* – $p \leq 0.01$ (Mann–Whitney–Wilcoxon's test, inductor alone vs. inductor + lipopeptide). Concentrations of CLPs used are indicated in Supplementary Table S1 (third column).

**Supplementary Table S3.** The characteristic time (*τ*) of calcein leakage upon the fusion of vesicles composed of DOPC/DOPG/CHOL (40/40/20 mol.%) and POPC/SM/CHOL (60/20/20 mol.%) mediated by 40 mM CaCl_2_ and 150 μM FP_SARS(816-827)_, respectively.

| | $\text{CaCl}_2$ | $\text{FP}_{\text{SARS}(816-827)}$ |
| --- | --- | --- |
| *lipopeptide* | $\tau, \text{min}$ | $\tau, \text{min}$ |
| – | $62 \pm 9$ | $15 \pm 4$ |
| *Aculeacin A* | $1 \pm 1^*$ | $35 \pm 6^*$ |
| *Anidulafungin* | $16 \pm 4^*$ | $15 \pm 7$ |
| *Caspofungin* | $6 \pm 1^*$ | $6 \pm 3$ |
| *Fengycin* | – | $14 \pm 5$ |
| *Iturin A* | $79 \pm 6$ | $90 \pm 25^*$ |
| *Mycosubtilin* | $7 \pm 4^*$ | $16 \pm 4$ |
| *Surfactin* | $26 \pm 7^*$ | $5 \pm 2$ |
| *Syringostatin A* | $52 \pm 33$ | $17 \pm 6$ |
| *Syringotoxin B* | $31 \pm 7^*$ | $6 \pm 3$ |
| *Daptomycin* | $18 \pm 9^*$ | $1 \pm 1$ |
| *Polymyxin B* | $82 \pm 38$ | $1 \pm 1$ |

\* – $p \leq 0.01$ (Mann–Whitney–Wilcoxon's test, inductor alone vs. inductor + lipopeptide) Concentrations of CLPs used are indicated in Supplementary Table S1 (third column).

**Supplementary Table S4.** The effect of the CLPs on the boundary potential (*φ_b_*) of membranes composed of POPC/SM/CHOL (60/20/20 mol.%).

| *lipopeptide* | $\Delta\phi_b, \text{mV}$ |
| --- | --- |
| *Aculeacin A* | $-68 \pm 23$ |
| *Anidulafungin* | $-2 \pm 6$ |
| *Caspofungin* | $-9 \pm 2$ |
| *Fengycin* | $-16 \pm 2$ |
| *Iturin A* | $-13 \pm 2$ |
| *Surfactin* | $-4 \pm 5$ |
| *Syringostatin A* | $-3 \pm 3$ |
| *Syringotoxin B* | $-6 \pm 2$ |
| *Daptomycin* | $-8 \pm 1$ |
| *Polymyxin B* | $9 \pm 4$ |

Concentrations of CLPs used are indicated in Supplementary Table S1 (third column).

**Supplementary Figure S1.**
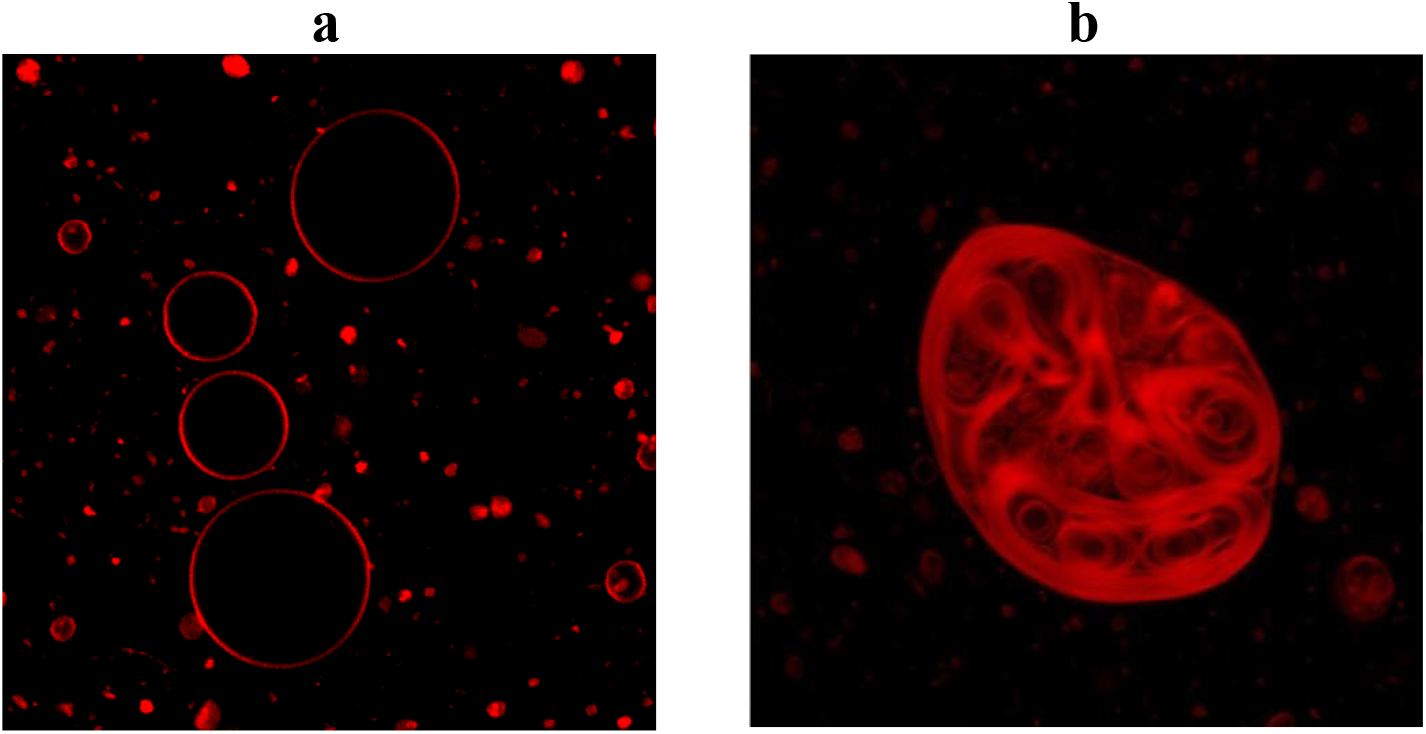
Typical fluorescence micrographs of giant unilamellar vesicles composed of POPC/SM/CHOL (60/20/20 mol%) and 1 mol.% of fluorescent lipid probe Rh-DPPE in the absence (**a**) and presence of 150 μM of the short fragment of SARS-CoV-2 fusion peptide 1 (816-SFIEDLLFNKVT-827, FP_SARS(816-827)_) (**b**). The size of each image is 50 μm × 50 μm.

**Supplementary Figure S2.**
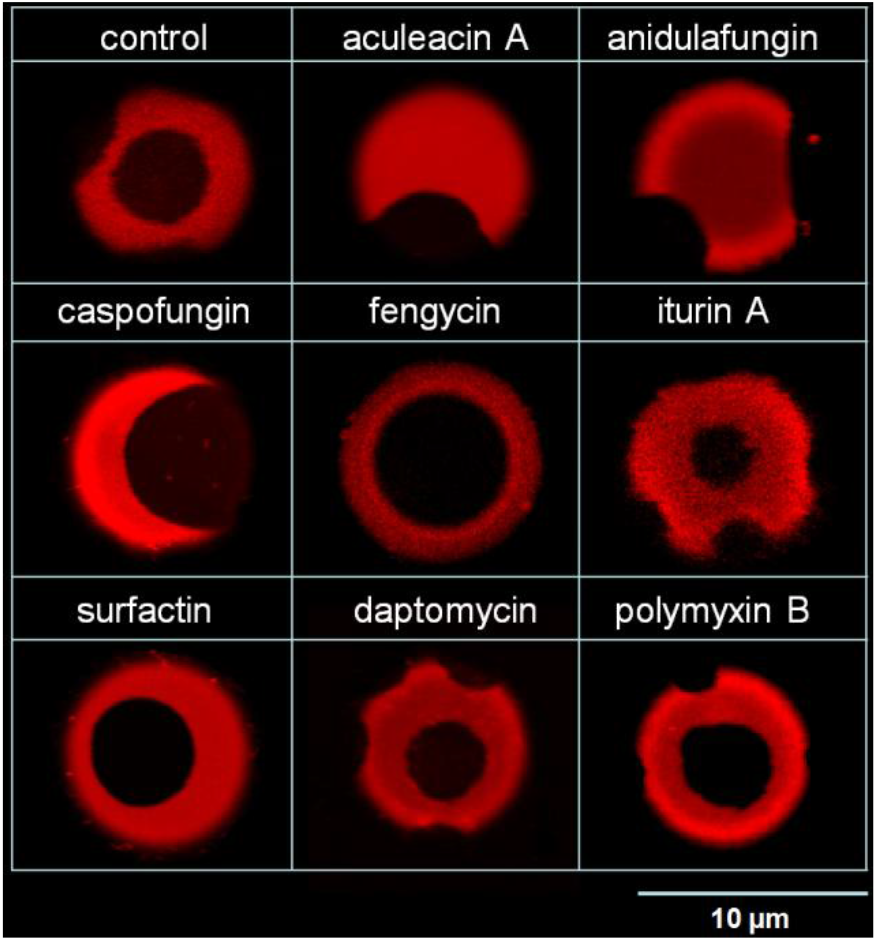
Typical fluorescence micrographs of giant unilamellar vesicles composed of DOPC/SM/CHOL (40/40/20 mol.%) and 1 mol.% of fluorescent lipid probe Rh-DPPE in the absence (control) and presence of different CLPs at a lipid:CLP molar ratio of 25:1.

